# Recurrent Independent Pseudogenization Events of the Sperm Fertilization Gene ZP3r in Apes and Monkeys

**DOI:** 10.1101/2021.10.15.464600

**Authors:** J.A. Carlisle, D.H. Gurbuz, W. J. Swanson

## Abstract

In mice, ZP3r/sp56 is a binding partner to the egg coat protein ZP3 and may mediate induction of the acrosome reaction. ZP3r, as a member of the RCA cluster, is surrounded by paralogs, some of which have been shown to be evolving under positive selection. Sequence divergence paired with paralogous relationships with neighboring genes, has complicated the accurate identification of the human ZP3r ortholog. Here, we phylogenetically and syntenically resolve that the human ortholog of ZP3r is the pseudogene C4BPAP1. We investigate the evolution of this gene within primates. We observe independent pseudogenization events of ZP3r in all Apes with the exception of Orangutans, and many monkey species. ZP3r in both primates that retain ZP3r and rodents contains positively selected sites. We hypothesize that redundant mechanisms mediate ZP3 recognition in mammals and ZP3r’s relative importance to ZP recognition varies across species.

## Introduction

Complex molecular interactions between the sperm and egg mediate fertilization (J. A. Carlisle & Swanson, 2020; Swanson & Vacquier, 2002). Although recent discoveries have described many important molecules mediating mammalian sperm-egg plasma membrane fusion, the molecular mediators of sperm-egg coat interactions remain ambiguous (J. A. Carlisle & Swanson, 2020). The glycoprotinaceous egg molecules ZP2 and ZP3 have been shown to bind sperm in a species-specific manner, indicating that these molecules may be involved in sperm-egg interactions (Avella, Baibakov, & Dean, 2014; Bleil & Wassarman, 1980; J. A. Carlisle & Swanson, 2020; Litscher, Williams, & Wassarman, 2009). While there is no known sperm protein binding partner of ZP2, ZP3r (formally known as sp56) is described as the receptor of ZP3 in mice (Buffone et al., 2008; Wassarman, 2009). ZP3r is a sperm acrosomal protein that becomes transiently exposed on the sperm head post-capacitation in mice (Muro, Buffone, Okabe, & Gerton, 2012). Isolated ZP3r inhibits sperm binding by binding mouse eggs *in vitro* and specifically binds ZP3 as shown by photoaffinity cross-linking (Bleil & Wassarman, 1990; Buffone et al., 2008). Despite these compelling results, mouse knockouts of ZP3r do not result in observable reductions in fertility, however this may be due to alternative assays being needed to observe ZP3r’s function (Adham, 1998; Muro et al., 2012; Okabe, 2018). For example, the sperm protein PKDREJ, while not causing infertility in male mice knockouts, does lead to a delay in the induction of the acrosome reaction by ZP recognition and a reduction in male fertility compared to wild type animals in sequential mating trials (Miyata et al., 2016; Sutton, Jungnickel, & Florman, 2008; Sutton, Jungnickel, Ward, Harris, & Florman, 2006). Multiple proteins, including ZP3r, may contribute to sperm recognition of the egg coat, and have redundant functions.

Identification of the human ortholog of ZP3r has been controversial. Previous studies have misidentified human ZP3r as either SELENBP1 or C4BPA, due to nomenclature confusion or difficulties in establishing orthology respectively (Morgan & Hart, 2019; Morgan, Loughran, Walsh, Harrison, & O’Connell, 2010, 2017). ZP3r is found in chromosome 1 amongst paralogous protein-coding genes that make up the RCA cluster (Hourcade, Holers, & Atkinson, 1989; Krushkal, Bat, & Gigli, 2000). Many of these genes are diverging rapidly between species (Hart et al., 2018). This sequence divergence of the paralogs can further complicate accurate ortholog identification. In this study we demonstrate using syntenic and phylogenetic analysis that C4BPAP1 is the primate ortholog of ZP3r. We examine the evolution of ZP3r in primates and uncover a pattern of recurrent independent pseudogenizations of ZP3r in Great Apes and Monkeys. This work highlights the complexities of identifying orthologs between species, particularly when pseudogenizations have occurred, and indicates that redundant mechanisms of gamete recognition may lead to the loss of reproductive genes.

## Results and Discussion

### C4BPAP1 is the ortholog of mouse ZP3r

Although well characterized in mice, the identification of primate ZP3r has been contentious. In mice, ZP3r is located within the RCA cluster. An examination of this genomic region in humans reveals the paralogs C4BPA and C4BPAP1. C4BPA is a large glycoprotein that acts as an inhibitor within the complement system (Okroj M., 2018). C4BPAP1 shares a domain structure and sequence similarity to C4BPA but contains a premature stop codon in Exon 2 that is fixed in humans and suggests pseudogenization. Human C4BPA and C4BPAP1 are composed of 11 exons which contain 8 CCP/sushi domains and a C-terminal transmembrane domain (Hofmeyer et al., 2013). Both C4BPAP1 and C4BPA contain an additional CCP domain to mouse ZP3r which is missing CCP domain 7. A recent investigation identified C4BPA as the human ortholog of ZP3r, perhaps overlooking C4BPAP1 since it is pseudogenized in humans (Morgan & Hart, 2019). The study hypothesized that a duplication in rodents of C4BP, the rodent ortholog of human C4BPA, led to the evolution of rodent ZP3r (Morgan & Hart, 2019). However, human C4BPA is known to function in immunity and its highest tissue expression is in the liver, inconsistent with a function as the sperm fertilization gene ZP3r (Carithers & Moore, 2015; Okroj M., 2018). Meanwhile, although pseudogenized, C4BPAP1 shows highest RNA expression in the testis, consistent with an ancestral function as a sperm fertilization gene (Carithers & Moore, 2015). Genes that have been recently pseudogenized are often still expressed until completely knocked out (Bekpen et al., 2009).

Using phylogenetic analysis and syntenic mapping we showed that C4BPAP1 is the human ortholog of mouse ZP3r (Figure 1). A protein alignment of *Homo sapiens* and *Macaca mulatta* C4BPAP1 and C4BPA and *Mus musculus* and *Rattus norvegicus* ZP3r and C4BP sequences were used to construct a maximum likelihood phylogeny. Primate C4BPAP1 and Rodent ZP3r clustered separately from Primate C4BPA and Rodent C4BP, indicating that Primate ZP3r (C4BPAP1), not C4BPA, is the ortholog of rodent ZP3r (Figure 1B). Further, we used the best reciprocal blast hits of ZP3r/C4BPAP1 (stop codon removed), C4BPA, and neighboring RCA cluster gene transcripts between the mouse and human genome to establish syntenic relationships. Syntenic comparison between the region of the human and mouse RCA clusters containing C4BPAP1 and ZP3r respectively, support C4BPAP1 as the human ZP3r ortholog. In humans, C4BPAP1 is located between C4BPA and CD55, as is ZP3r in rodents (Figure 1A) (Kent et al., 2002).

**Figure 1:**
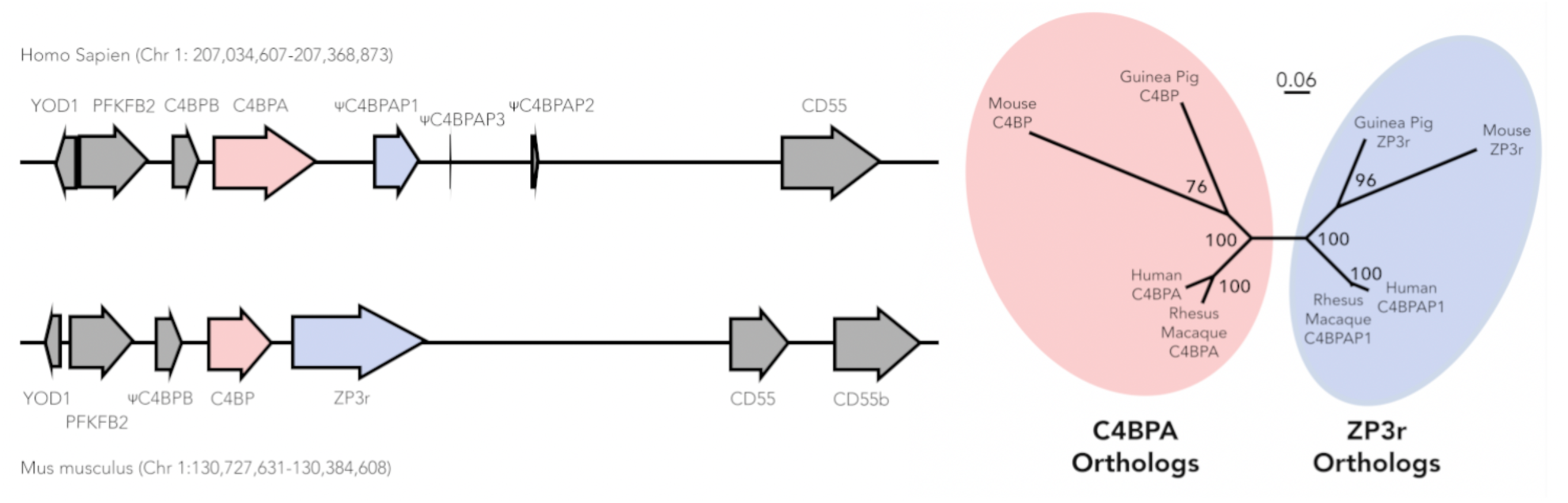
Syntenic and phylogenetic analysis indicates that C4BPAP1 is the human ortholog of mouse ZP3r. A.) Syntenic comparison of the genomic region between Mus musculus and Homo sapiens reveals that C4BPAP1 is syntenic to mouse ZP3r. B) A protein alignment of C4BPAP1 and C4BPA from *Homo sapiens* and *Macaca mulatta* and ZP3r and C4BP from *Mus musculus* and *Rattus norvegicus* was constructed using Clustal Omega. This protein alignment was used to construct a maximum likelihood phylogeny with bootstrapping. In the phylogeny, Primate C4BPAP1 and Rodent ZP3r cluster separately from Primate C4BPA and Rodent C4BP, indicating that Primate ZP3r (C4BPAP1), not C4BPA, is the ortholog of rodent ZP3r. The phylogenetic inference tool RAxML-NG was used to construct the phylogenetic tree with the LG substitution matrix. RaxML-NG conducts maximum likelihood based phylogenetic inference and provides branch support using non-parametric bootstrapping. The best scoring topology of 20 starting trees (10 random and 10 parsimony-based) was chosen. RaxML-NG was used to perform non-parametric bootstrapping with 1000 re-samplings that were used to re-infer a tree for each bootstrap replicate MSA.

Previous research identified elevated linkage disequilibrium between the region of the human genome containing ZP3r and the region containing ZP3 suggestive of coevolution between these loci (Rohlfs, Swanson, & Weir, 2010). Since ZP3r is pseudogenized in humans, this was possibly a false positive result or a complex association. An alternative hypothesis would be that the human sperm receptor for ZP3 is located nearby the pseudogenized human ZP3r. However, none of the annotated genes within the region shown to be in LD with ZP3 show testes-specific expression (Carithers & Moore, 2015).

### ZP3r has been repeatedly and rapidly pseudogenized in Apes

C4BPA and ZP3r/C4BPAP1 are members of the RCA cluster, the genes in this locus are largely conserved across even distantly related species, with sequence variation between species being driven by positive selection, indels, and intragenic domain duplications and losses (Garcia-Fernandez, Vilches-Arroyo, Olavarrieta, Perez-Perez, & Rodriguez de Cordoba, 2021; Heinen et al., 2006; Sanchez-Corral, Pardo-Manuel de Villena, Rey-Campos, & Rodriguez de Cordoba, 1993; Wu, Li, & Zhang, 2012). However, some variation in RCA cluster gene content driven by clade-specific duplication or loss events have also been observed (Pardo-Manuel de Villena, 1995; Sanchez-Corral et al., 1993; Wu et al., 2012). Notably, C4BPB is pseudogenized in mice and there is evidence of two additional pseudogenized duplications of C4BPA found in humans (C4BPAP2 and C4BPAP3) (Kent et al., 2002; Pardo-Manuel de Villena, 1995). However, primate ZP3r/C4BPAP1 is unique in independently acquiring pseudogenization events in most apes and several monkey species (Figure 2). Although there are examples in the literature of repeated pseudogenization events of genes across species, it is rare for independent events to occur within a closely related clade (Bainova et al., 2014; Velova, Gutowska-Ding, Burt, & Vinkler, 2018). Remarkably, since the common ancestor of all apes (∼16-20 mya), at least four unique pseudogenization events of ZP3r have occurred (Figure 2) (Chatterjee, Ho, Barnes, & Groves, 2009).

**Figure 2:**
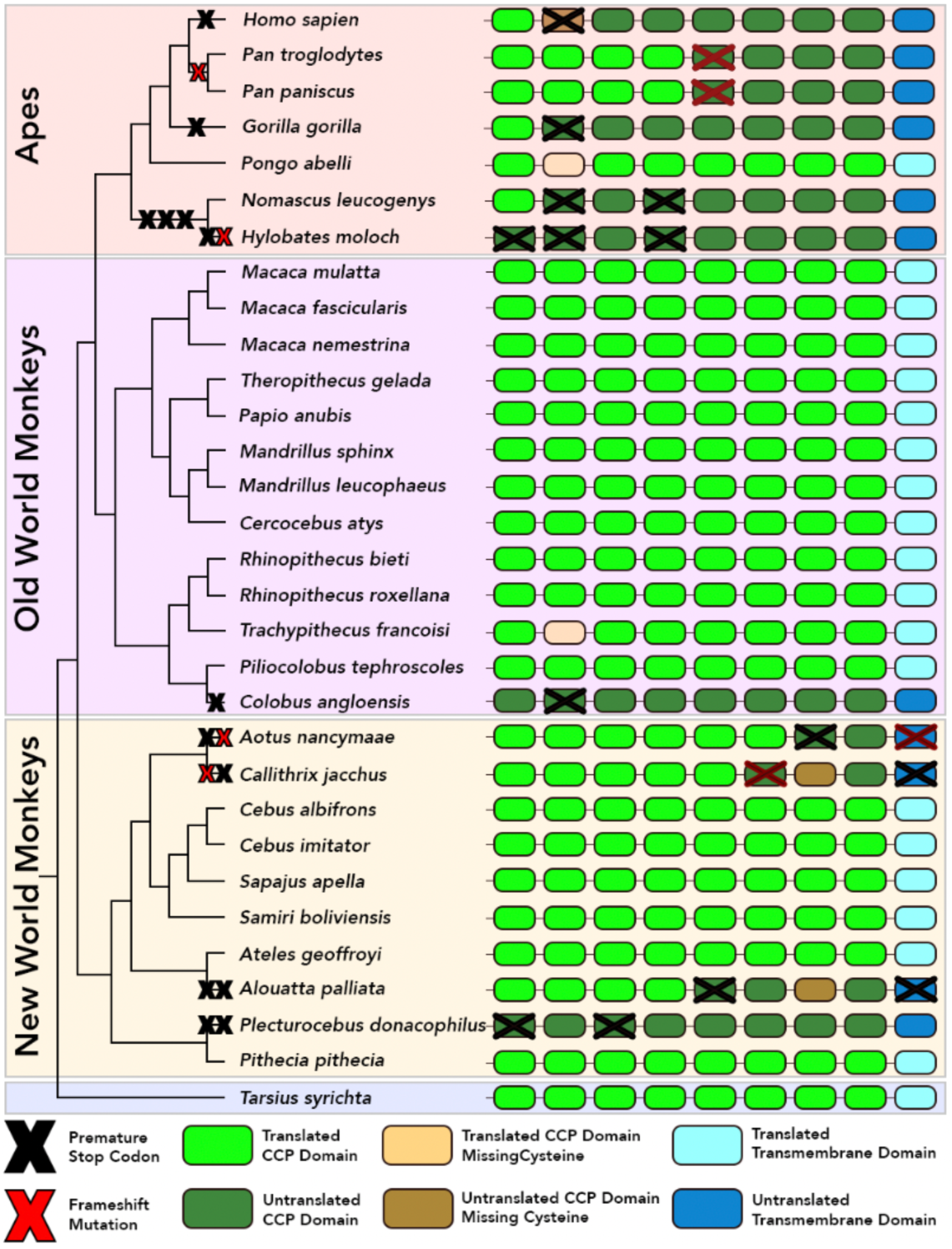
Recurrent and independent pseudogenization events of ZP3r in apes and monkeys. We predicted the exonic sequences of ZP3r from primate genomes using the Protein2Genome command of the program Exonerate version 2.2.0 (Slater & Birney, 2005). The top scoring prediction from Exonerate was used to define the paralog’s exons. We used human C4BPAP1, with the stop codon in CCP2 removed, as the protein query in Exonerate. Using HMMER, we identified CCP domains in the ZP3r sequences. On the right side is a not-to-scale cartoon of the CCP domains (Green ovals) and C-terminal transmembrane domain (Blue ovals) found in ZP3r in each species. The loss of a CCP domain caused by a missing structurally important cysteine are shown as Yellow ovals. Red crosses indicate an insertion/deletion mutation causing a frameshift mutation; Black crosses indicate a premature stop codon causing pseudogenization. Darker colored CCP domains indicate regions of ZP3r that would not be translated due to a pseudogenization event. Within some CCP domains, multiple mutations have occurred. See Supplementary Figure 1 for a protein alignment of Ape ZP3r with pseudogenizing mutations marked. The cladogram on the left has pseudogenization events marked. For branches with multiple mutations the order in which these mutations occurred are unknown.

Parsimony analysis of the pseudogenized ZP3r sequences indicate 9 independent pseudogenization events have occurred in primates. Remarkably, many of these pseudogenizing mutations occurred independently in closely related species and are located in distinct codons (Supplementary Figure 1). With the exception of Orangutan (*Pongo abelli*), C4BPAP1 has been pseudogenized in all apes (Human, Chimpanzee, Bonobo, Gorilla, Gibbons) (Figure 2). This rapid, repeated pseudogenization appears to be an extreme example of gene loss in apes. Gorillas, Humans, and Gibbons all have premature stop codons within CCP domain 2, all in different codons (Supplementary Figure 1). Orangutan’s ZP3r does not have any pseudogenizing mutations, however, its second CCP domain is missing a conserved and potentially structurally important cysteine that may disrupt the overall structure of the protein. In 10 New World monkey (NWM), 13 Old World monkey (OWM), and one Tarsier genome assemblies, we identified the full ZP3r locus. Out of the 10 NWM genomes examined, 4 contained pseudogenizing mutations unique to that NWM species (Figure 2). In OWMs, only one species, *Colobus angoloensis*, had a pseudogenizing mutation within ZP3r (Figure 2).

Recurrent, lineage-specific gene loss events between closely related species is suggestive of strong selection for gene loss. There is no obvious correlation between ZP3 sequence and glycosylation state and ZP3r loss in primates, therefore, it is still unclear what is driving the loss of ZP3r in primates. Phylogenetic analysis of all individual CCP domains found in human C4BPA and C4BPAP1 and mouse C4BP and ZP3r indicate no evidence of concerted evolution between or within genes that could explain the repeated pseudogenization events (Supplementary Figure 2). Further, a search of the human genome reveals no new duplications of ZP3r that could be fulfilling its receptor function. However, a more distantly related paralog with low sequence similarity could be performing ZP3r’s function. Protein structure changes more slowly than protein sequence, therefore a paralog with low sequence identity may still retain similar function.

### ZP3r Evolves Under Positive Selection

A recurrent feature of gamete recognition proteins are signatures of positive selection, potentially created through sexual selection or sexual conflict between the sperm and the egg (J. A. Carlisle & Swanson, 2020). Genes mediating immune system functions are also frequently undergoing positive selection due to host-pathogen interactions driving arms race dynamics (Lazzaro, 2012). So, it is unsurprising that both ZP3 and C4BPA have both been shown in previous studies to be undergoing positive selection in rodents and primates (Hart et al., 2018; Morgan & Hart, 2019; Rohlfs et al., 2010; Swann, Cooper, & Breed, 2007; Swann, Cooper, & Breed, 2017; Swanson, Yang, Wolfner, & Aquadro, 2001). In this study, we estimated values of *d*_*N*_*/d*_*S*_ for ZP3r, C4BPA, and ZP3 in rodents and primates using the codeml program of PAML 4.8 (Yang, 1997, 2007). For primate ZP3r, we only analyzed full coding sequences, no pseudogenized primate sequences were included. We compared models of selection using a likelihood ratio test (LRT) between neutral models and models with positive selection. Specifically, we compared M1 v. M2, M7 v. M8, and M8a v. M8 (Swanson, Nielsen, & Yang, 2003; Yang, Nielsen, Goldman, & Pedersen, 2000).

We detected positively selected sites in ZP3r, C4BP, and ZP3 in rodents and ZP3r and C4BPA in primates, using the M8a v M8 comparison (Table 1). Signatures of positive selection in rodent and primate ZP3r is suggestive of functionally important genetic innovation being selected for within both clades. Because interacting reproductive proteins must co-evolve to maintain reproductive compatibility, ZP3r’s rapid evolution could be driven by the evolution of its putative binding partner ZP3. Although positively selected sites were not detected in primate ZP3 in this study, previous population genetic analysis has detected selection on ZP3 in humans (Hart et al., 2018; Rohlfs et al., 2010). Since ZP3r is undergoing positive selection in primates, ZP3r’s repeated and independent pseudogenization in primates is likely not driven by relaxed selection.

**Table 1:** ZP3r and C4BPA contain positively selected sites in rodents and primates. Codon substitution models were used to analyze sequences of ZP3, ZP3r/C4BPAP1, C4BP/C4BPA in rodents and primates. Site models allowing for several neutral models (M1a, M7, and M8a) or selection models (M2a, M8, and M8a) allowing for variation among sites, were fit to the data using PAML. In this table are the results from M8a v M8 comparison, for the results from other model comparisons see Supplementary Table 1. In rodents, sites under positive selection were detected in ZP3, ZP3r, and C4BP. In primates, M8a v M8 model comparison indicated sites under positive selection were detected in ZP3r and C4BPA, but not ZP3. Estimates of the likelihood ratio statistic (**-2Δl)**, *d*_*N*_*/d*_*S*_, and the percentage of sites that are under positive selection are given. (*, significant at P < 0.05; **, significant at P < 0.005.)

| Gene | Clade | Model | -2ΔI | dN/dS | % Positively Selected Sites |
| --- | --- | --- | --- | --- | --- |
| ZP3 | Rodents | M8 v M8a | **14.63 | 3.66585 | 1.6 |
| ZP3 | Primates | M8 v M8a | 1.75 |  |  |
| ZP3r | Rodents | M8 v M8a | **84.56 | 3.12174 | 7.5 |
| ZP3r | Primates | M8 v M8a | *3.89 | 3.01771 | 3.4 |
| C4BP | Rodents | M8 v M8a | **100.18 | 3.08952 | 9.7 |
| C4BPA | Primates | M8 v M8a | **86.11 | 3.62581 | 12.6 |

## Conclusion

Despite their functional importance, the molecular mediators of fertilization have been poorly described in mammals, particularly for identifying sperm proteins mediating egg coat recognition (J. A. Carlisle & Swanson, 2020). Difficulty in finding sperm receptors to egg coat proteins may be driven by functional redundancy causing many fertilization genes to be nonessential contributors to gamete recognition. Typically, protein functional redundancy refers to paralogous proteins that are structurally similar, that maintain the same interaction partners, and whose loss can be compensated for by their paralog. However, proteins can also be functionally redundant without being paralogous or structurally similar. For example, the acrosomal sperm proteins Zona Pellucida Binding Protein (ZPBP/sp38) and acrosin are structurally unrelated proteins that competitively interact with the ZP in boars (Lin, Roy, Yan, Burns, & Matzuk, 2007; Mori, Baba, Iwamatsu, & Mori, 1993). Functional redundancy of genes mediating fertilization could lead to clade-specific gene loss events or changes in relative functional importance between species. Again, reflecting on acrosin, knockouts of acrosin in mice (Mus musculus) result in infertility; yet, in hamsters (*Mesocricetus auratus*) acrosin is essential for zona penetration (Baba, Azuma, Kashiwabara, & Toyoda, 1994; Hirose et al., 2020). Together, these results indicate that functional redundancy between ZPBP, acrosin, and potentially other unknown sperm proteins, allow the relative importance of acrosin to sperm bypassing the ZP to vary between species.

In this study, we demonstrate that the testes-expressed pseudogene C4BPAP1 is the human ortholog of rodent ZP3r using phylogenetic and syntenic analysis. While ZP3r is associated with ZP binding in mice, ZP3r shows repeated pseudogenization in primates (at least 9 times), most notably in apes. Recurrent independent pseudogenizations of a rapidly evolving protein are rarely discussed in the literature, and their existence is surprising. While usually rapid divergence is focused on sequence diversification, changes in gene content caused by gene gains and loss events could also be a significant contributor to molecular diversity and tolerated due to functional redundancy (J. A. Carlisle, Glenski, M.A., Swanson, W.J., 2021). ZP3r is a nonessential fertilization gene in mice, who may be one of many proteins interacting with ZP3 (Miyata et al., 2016; Muro et al., 2012; Okabe, 2018). ZP3r’s repeated loss in many primates, particularly apes, despite being subject to positive selection in other primate species, indicates that the relevant importance of ZP3r to fertilization differs across primates. This difference could be due to the emergence or increase in relative importance of a different fertilization gene mediating ZP3 binding in primates. Differences in relative functional importance between clades may also partially explain why reproductive proteins are rapidly evolving in some clades and not others (J. A. Carlisle & Swanson, 2020). This study highlights the potential variability of molecular mechanisms of fertilization even within mammals and emphasizes the value of using diverse model systems for investigating mechanisms of fertilization.

## Contributions

JAC and WJS designed the research. JAC and DHG performed the research. JAC wrote the paper.

## Acknowledgements

This study was supported by NIH Grant HD076862 to Willie J. Swanson and a National Science Foundation Graduate Research Fellowship to Jolie A. Carlisle. Thank you to Dr. Damien Wilburn, Dr. Jan Aagaard, and Alberto Rivera for useful feedback. All research was performed on the traditional lands of the Duwamish Tribe. To learn more about the Duwamish Tribe and their continuing legacy, please visit https://www.duwamishtribe.org/.

---

**Supplementary Figure 1:**
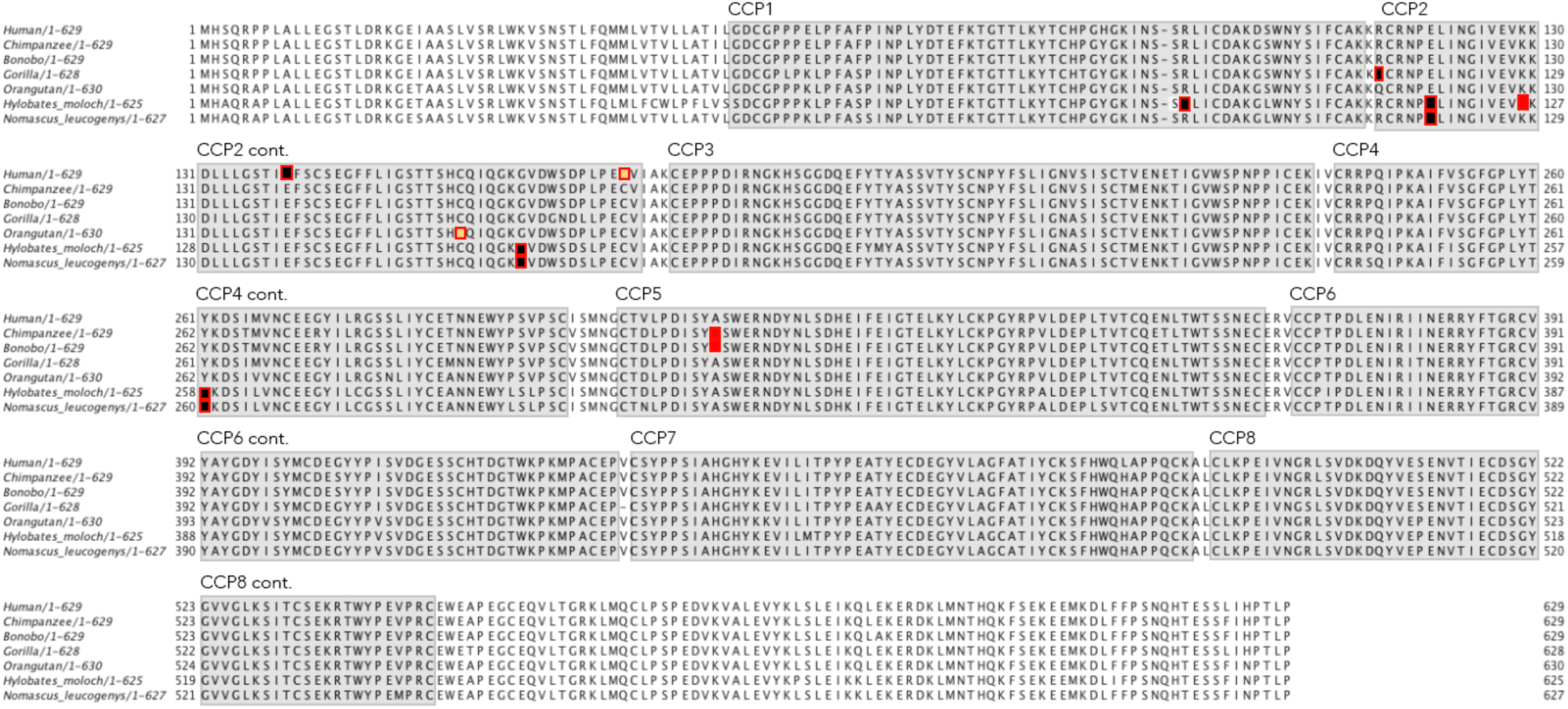
Protein phylogeny of rodent C4BPA and ZP3r CCP domains reveals no evidence of concerted evolution. Protein Alignment of C4BPAP1/ZP3r sequences in Apes. ZP3r in all apes, with the exception of Orangutans, have at least one mutation that results in premature termination of translation. Black squares indicate a premature stop codon; Red squares indicate an insertion/deletion mutation that results in premature termination of translation. Yellow squares indicate a cysteine that is structurally important to the CCP domain has been lost.

**Supplementary Figure 2:**
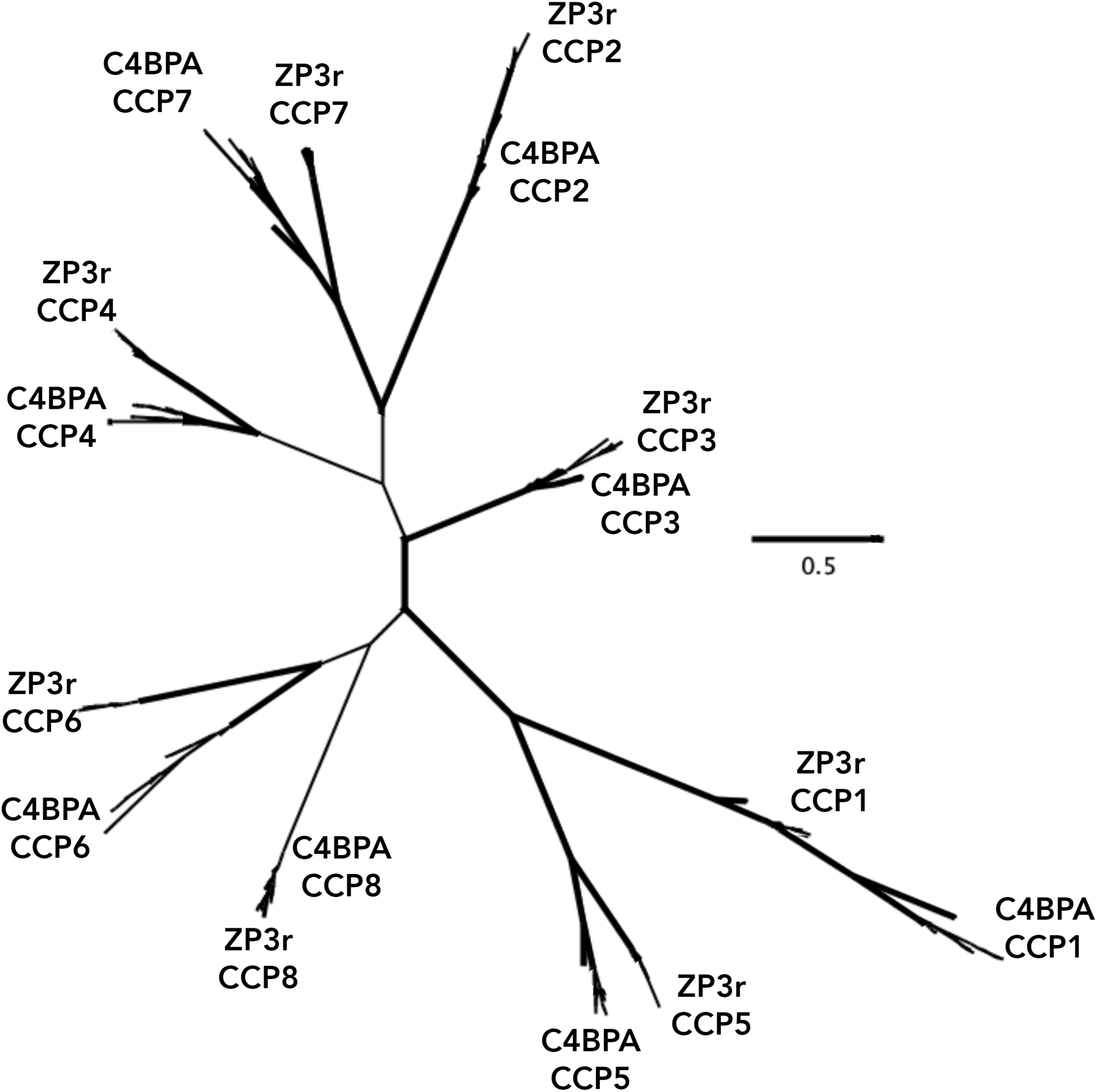
Protein phylogeny of primate C4BPA and ZP3r CCP domains reveals no evidence of concerted evolution. Bootstrap support greater than 70 is shown by a bold line. The separation of CCP domains between either C4BPA and ZP3r can be difficult to see in this image. See the attached newick tree file (Supplementary File 1) to more closely examine the phylogenetic relationships between protein domains.

**Supplementary Table 1:** ZP3r, C4BPA, and ZP3 all contain positively selected sites. Codon substitution models were used to analyze sequences of ZP3, ZP3r/C4BPAP1, C4BP/C4BPA in rodents and primates. Site models allowing for several neutral models (M1a, M7, and M8a) or selection models (M2a, M8, and M8a) allowing for variation among sites, were fit to the data using PAML. In rodents, sites under positive selection were detected in ZP3, ZP3r, and C4BP for all model comparisons. In primates, sites under positive selection were also detected in all three genes, but not for all model comparisons. Primate ZP3 only showed positively selected sites in a M7 v M8 comparison. A more powerful test (M8a v M8) did not detect positive selection in primate ZP3. Estimates of the likelihood ratio statistic (**-2Δl)**, *d*_*N*_*/d*_*S*_, and the percentage of sites that are under positive selection are given. (*, significant at P < 0.05; **, significant at P < 0.005.)

| Gene | Clade | Model | -2Δl | dN/dS | % Positively Selected Sites |
| --- | --- | --- | --- | --- | --- |
| ZP3 | Rodents | M1a v M2a | **11.16 | 5.41816 | 1.1 |
| ZP3 | Rodents | M7 v M8 | **17.61 | 3.66585 | 1.6 |
| ZP3 | Rodents | M8 v M8a | **14.63 | 3.66585 | 1.6 |
| ZP3 | Primates | M1a v M2a | 1.65 |  |  |
| ZP3 | Primates | M7 v M8 | *9.97 | 1.32 | 14.4 |
| ZP3 | Primates | M8 v M8a | 1.75 |  |  |
| ZP3r | Rodents | M1a v M2a | **101.47 | 4.19662 | 5.3 |
| ZP3r | Rodents | M7 v M8 | **109.43 | 3.12174 | 7.5 |
| ZP3r | Rodents | M8 v M8a | **84.56 | 3.12174 | 7.5 |
| ZP3r | Primates | M1a v M2a | 4.38 |  |  |
| ZP3r | Primates | M7 v M8 | *6.32 | 3.01771 | 3.4 |
| ZP3r | Primates | M8 v M8a | *3.89 | 3.01771 | 3.4 |
| C4BP | Rodents | M1a v M2a | **107.47 | 3.56697 | 8.1 |
| C4BP | Rodents | M7 v M8 | **115.69 | 3.08952 | 9.7 |
| C4BP | Rodents | M8 v M8a | **100.18 | 3.08952 | 9.7 |
| C4BPA | Primates | M1a v M2a | **86.48 | 3.72082 | 11.7 |
| C4BPA | Primates | M7 v M8 | **89.92 | 3.62581 | 12.6 |
| C4BPA | Primates | M8 v M8a | **86.11 | 3.62581 | 12.6 |

